# Newly unveiled meiosis elucidates the unreduced gamete frequency and its impact on evolution of the *Lemna minor* complex

**DOI:** 10.64898/2026.02.11.705290

**Authors:** Yuri Lee, Veit Schubert, Anton Stepanenko, Gihwan Kim, Luca Braglia, Ingo Schubert, Laura Morello

**Affiliations:** Institute of Agricultural Biology and Biotechnology, National Research Council (CNR-IBBA), Via Alfonso Corti 12, 20133 Milan, Italy; Leibniz Institute of Plant Genetics and Crop Plant Research (IPK), Gatersleben, 06466 Seeland, Germany; Department of Molecular Genetics, Institute of Cell Biology and Genetic Engineering, NASU, 03143 Kyiv, Ukraine

**Author notes:** author for correspondence: Yuri Lee. Further Email addresses: VS, AS, GK, LB, IS, LM.

**Keywords:** Duckweeds, meiosis, interspecific hybrids, *Lemna* section *Lemna*, sexual reproduction, unreduced male gametes

## Abstract

Fusion of gametes possessing meiotically reduced (haploid) chromosome complements is the main pathway of propagation among eukaryotes. However, duckweeds, the smallest angiosperms, propagate mainly vegetatively, and meiosis has not yet been documented in detail for this plant family. The more surprising was the recent evidence of rather frequent interspecific hybrids and triploid clonal accessions which became obvious by genome size measurements, genomic in situ hybridization (GISH) and combined plastid and nuclear DNA markers. These observations indicated sexual propagation involving reduced as well as unreduced male and female gametes in *Lemna minor* and *L. turionifera* leading to allodiploid and allotriploid hybrids (MT, MMT, MTT) and autotriploid *L. minor* (MMM) accessions. Here, we i) documented the meiotic stages of *Lemna* species for the first time; ii) provided evidence of unreduced male gametes through fluorescent in situ hybridization (FISH) with single locus probes; iii) determined their abundance in different individuals and iv) hypothesized about the reasons of unreduced male gamete formation. These findings open new insights into the modes of sexual reproduction and evolution of duckweeds which may be useful for future breeding efforts in this emerging crop.

## Introduction

Meiosis as a reductive cell division is essential for sexual reproduction. It comprises a single round of DNA replication followed by two cell divisions, separating homologous chromosomes during the first, and sister chromatids during the second division, thus generating cells with half of the chromosomes of the somatic mother cell. However, meiotic defects can produce viable gametes retaining somatic chromosome numbers, potentially leading to polyploid progeny (Brownfield and Köhler 2011; Zamariola et al. 2014). Plant polyploidy is widespread in natural ecosystems and in many major crops (Brownfield and Köhler, 2011). Therefore, understanding the cytological events underlying normal meiosis as well as the formation of unreduced gametes is fundamental for elucidation of evolutionary processes and provides a basis for plant breeding.

Duckweeds, the smallest flowering angiosperms, are an aquatic monocot family (Lemnaceae Martinov) which mainly reproduces through fast asexual propagation. *Lemna* L., the largest of five duckweed genera, comprises four sections (*Alatae* Hegelm., *Biformes* Landolt, *Lemna* L., and *Uninerves* Hegelm.) including so far 11 species and three interspecific hybrids as well as ploidy variants and backcrosses (for review see Morello et al. 2026). Even though these plants have been studied for over two centuries, due to their small size (0.6−9 mm in diameter), simple organismic structure (leaf-like fronds with one root), and rare flowering under natural and/or experimental condition, their taxonomy and phylogeny remained unclear till recently.

Recent applications of advanced molecular and cytological approaches have revealed surprisingly frequent auto-and allopolyploids which require sexual propagation (Braglia et al. 2024; Ernst et al. 2025; Stepanenko et al. 2025; Michael et al. 2025). Notably, the representative interspecific hybrid in sect. *Lemna, Lemna* × *japonica* Landolt, results from unidirectional crosses between *L. minor* L. (M) ♀ and *L. turionifera* Landolt (T) ♂ and exhibits variation in ploidy and genome composition (MT, MMT, MTT) (Braglia et al. 2021; Ernst et al. 2025), implying recurrent hybridization and the occurrence of unreduced gametes in both parental species. Consistent with this inference, naturally occurring autotriploid *L. minor* accessions have also been reported (Michael et al. 2026). However, direct experimental confirmation of unreduced gamete formation has been lacking so far.

In most flowering plants, interploidy crosses via fusion of reduced and unreduced gametes usually result in abnormal endosperm development and seed abortion, called “triploid block” (Cooper and Brink 1945; Köhler et al. 2021; Marks 1966). Duckweeds, can overcome this reproductive barrier, at least in part, possibly through the loss of genes of RNA-dependent DNA methylation (RdDM) pathways (Ernst et al. 2025). Their predominant clonal growth facilitates the persistence of triploid lineages and their sympatric diversification, despite sterility (Lee et al. 2025). Consequently, knowledge about the frequency of reduced and unreduced gametes in duckweeds is instrumental for elucidating their evolutionary dynamics.

Due to many small chromosomes and only few meristematic cells, reliable cytological studies are scarce for duckweeds (e.g., Cao et al. 2016; Hoang and Schubert 2017; Hoang et al. 2019, 2020, 2022; Stepanenko et al. 2025). Moreover, the rarity of natural flowering, the difficulty of inducing flower formation under experimental conditions and the smallness of flowers (flower <1 mm with one pistil and asynchronously growing two anthers surrounded by a spathe; see Fig. S1 in Lee et al. 2025) prevented detailed studies of meiosis, despite the obvious need for assessing the evolutionary prospects of duckweeds. We are aware of only a very limited number of studies mentioning duckweed meiosis (Caldwell 1899; Maheshwari and Kapil 1963) without photo-documentation.

This study provides the first detailed cytological characterization of male meiotic stages, a protocol for their preparation from young flowers and for analyzing the diverse final meiotic products in the duckweed section *Lemna*. By examining *L. minor* and *L. turionifera*, we quantified the meiotic products of different ploidy levels, assessing the occurrence and frequency of reduced and unreduced gametes in these predominantly clonally propagating plants.

## Results

### Microscopic observation and determination of the male meiotic stages

Pollen mother cells at the zygotene stage of *L. turionifera* 9434 and *L. minor* 5500, were first observed in anthers of minimum sizes, i.e. 143 × 134 μm and 157 × 119 μm (L × W), respectively (Fig. 1). Pollen mother cells of the two *Lemna* species stained with 4′,6-diamidino-2-phenylindole (DAPI) displayed the typical meiotic stages, such as zygotene, pachytene, metaphase I and II, similar to those of other eukaryotes (Fig. 2). Metaphase I of *L. turionifera* 9434 and 7432 showed the presence of 21 bivalents, indicating a haploid chromosome number relative to the somatic complement. Meiosis was regular in both species and the distinct meiotic stages appeared synchronously.

**Figure 1.**
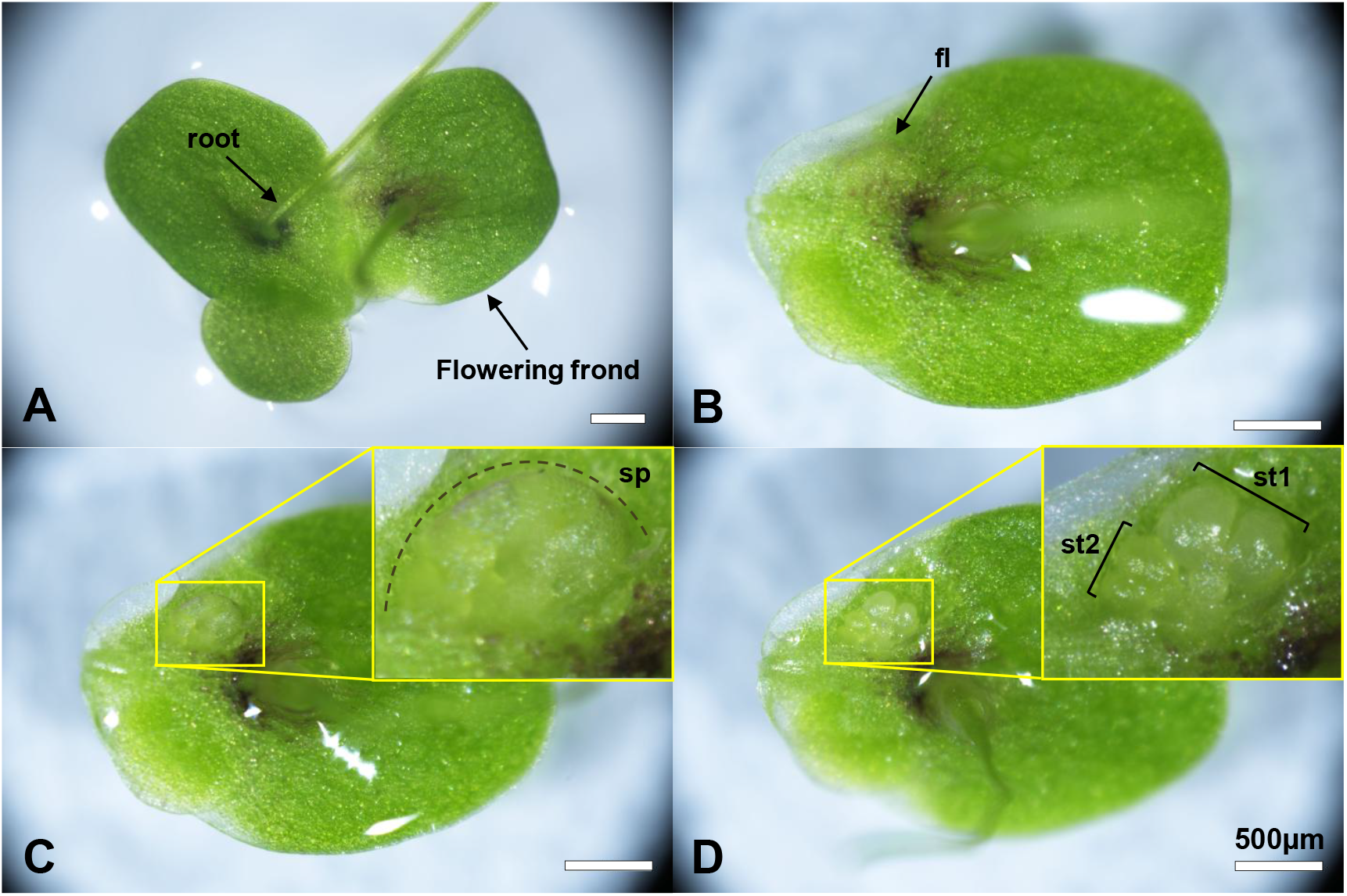
Different preparation stages of one individual of *Lemna turionifera* clone 7432 bearing immature flower (A) abaxial side of three cohering fronds, one of which is flowering; (B) the detached flowering (fl) frond; (C) the young flower surrounded by the reddish spathe (sp) is visible after removing frond’s abaxial surface; (D) the young, developing primary stamen (st1) and the secondary stamen (st2) are visible after removing the spathe.

**Figure 2.**
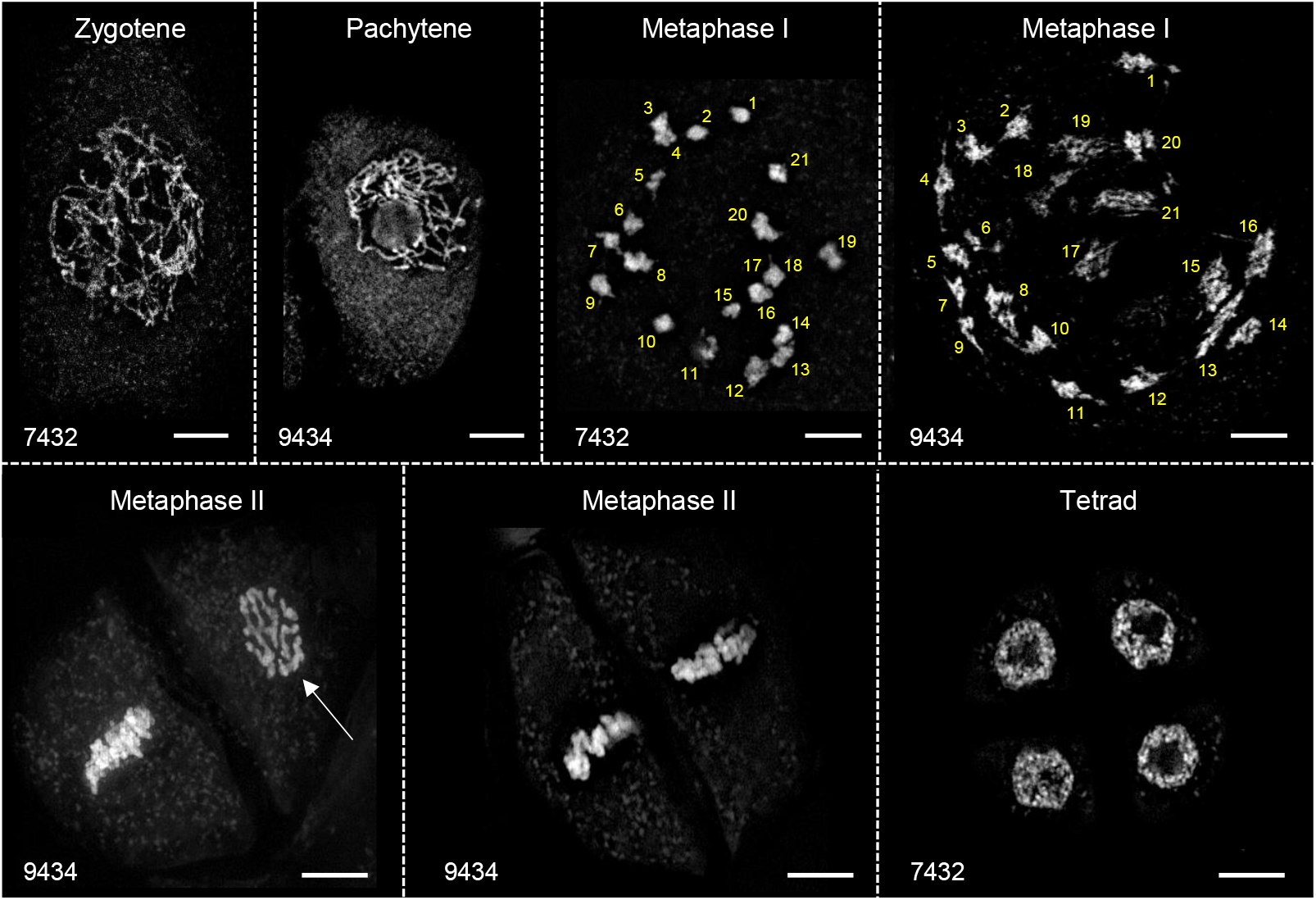
Meiotic stages of *Lemna turionifera* (accessions 9434 and 7432). Metaphase I contains 21 bivalents in both accessions. The right cell of the two different metaphase II meiocytes shows the chromosomes from top (arrow). Bars = 5µm

Tetrads as the meiotic end-product were detected in developing anthers with mean sizes of 175 × 144 μm for *L. turionifera* 9434, 196 × 153 μm for *L. turionifera* 7432, and 219 × 181 μm for *L. minor* 5500. Notably, the anther sizes corresponding to the tetrad stage varied among the three accessions, suggesting that anther size alone cannot serve as an absolute criterion for identifying the developmental stage, even within the same species. At the tetrad stage of the primary anther, the secondary anther typically exhibited cells at the zygotene to metaphase I stages.

### Detection of various meiotic products and confirmation of nuclei number in each cell

At the last stage of meiosis, across all three clones, most tetrads displayed a tetragonal-like arrangement of the resulting four cells. Flat clover leaf- and T-shaped forms were also observed, albeit at a lower frequency. In addition to tetrads, dyads and triads were also detected. In total, 4710 cells were observed. *L. turionifera* 9434 and 7432 produced 96.9% and 98.8% reduced gametes as well as 3.1% and 1.2% unreduced gametes, respectively. *L. minor* 5500 showed 96.3% reduced and 3.7% unreduced gametes (Table 1). The frequency of different meiotic end-products varied not only between accessions but also among individuals within the same accession.

**Table 1.** Numbers and ratio of reduced (1n) and unreduced (2n) gametes derived from diverse meiotic products (dyads, triads, and tetrads) in individual anthers of three *Lemna* accessions.

|  | 2n gamete<br>number<br>from dyad | 2n gamete<br>number<br>from triad | 1n gamete<br>number<br>from triad | 1n gamete<br>number<br>from tetrad | 2n gamete<br>rate (%) | 1n gamete<br>rate (%) |
| --- | --- | --- | --- | --- | --- | --- |
| *Lemna turionifera* 9434 | 40 | 0 | 0 | 544 | 6.85 | 93.15 |
|  | 2 | 0 | 0 | 440 | 0.45 | 99.55 |
|  | 12 | 2 | 4 | 348 | 3.83 | 96.17 |
|  | 0 | 0 | 0 | 404 | 0.00 | 100.00 |
| total | 54 | 2 | 4 | 1736 | 3.12 | 96.88 |
| *L. turionifera* 7432 | 2 | 2 | 4 | 420 | 0.93 | 99.07 |
|  | 4 | 0 | 0 | 556 | 0.71 | 99.29 |
|  | 0 | 0 | 0 | 112 | 0.00 | 100.00 |
|  | 4 | 0 | 0 | 580 | 0.68 | 99.32 |
|  | 0 | 0 | 0 | 412 | 0.00 | 100.00 |
|  | 4 | 2 | 4 | 404 | 1.45 | 98.55 |
|  | 6 | 4 | 8 | 188 | 4.85 | 95.15 |
| total | 14 | 6 | 12 | 1584 | 1.24 | 98.76 |
| *L. minor* 5500 | 2 | 0 | 0 | 372 | 0.53 | 99.47 |
|  | 4 | 1 | 2 | 200 | 2.42 | 97.58 |
|  | 2 | 2 | 4 | 268 | 1.45 | 98.55 |
|  | 32 | 11 | 22 | 292 | 12.04 | 87.96 |
|  | 6 | 0 | 0 | 336 | 1.75 | 98.25 |
|  | 0 | 0 | 0 | 452 | 0.00 | 100.00 |
| total | 40 | 13 | 26 | 1348 | 3.71 | 96.29 |

Cells of dyads and their nuclei were noticeably larger than those of tetrads (Fig. S1; Videos S1, S2). Fluorescence in situ hybridization (FISH) revealed the same number of rDNA signals in dyads within their two (or already fused) nuclei as observed in somatic cell nuclei. On the other hand, each tetrad cell nucleus showed only one 45S rDNA signal (Fig. 3). This was to be expected for single loci of 5S and 45S rDNA, as reported for *Lemna* species so far (Hoang et al. 2019). Therefore, the presence of tetrads, dyads and a few triads is consistent with the occurrence of reduced and unreduced gametes, respectively, in the three clones of the two frequently hybridizing *Lemna* species.

**Figure 3.**
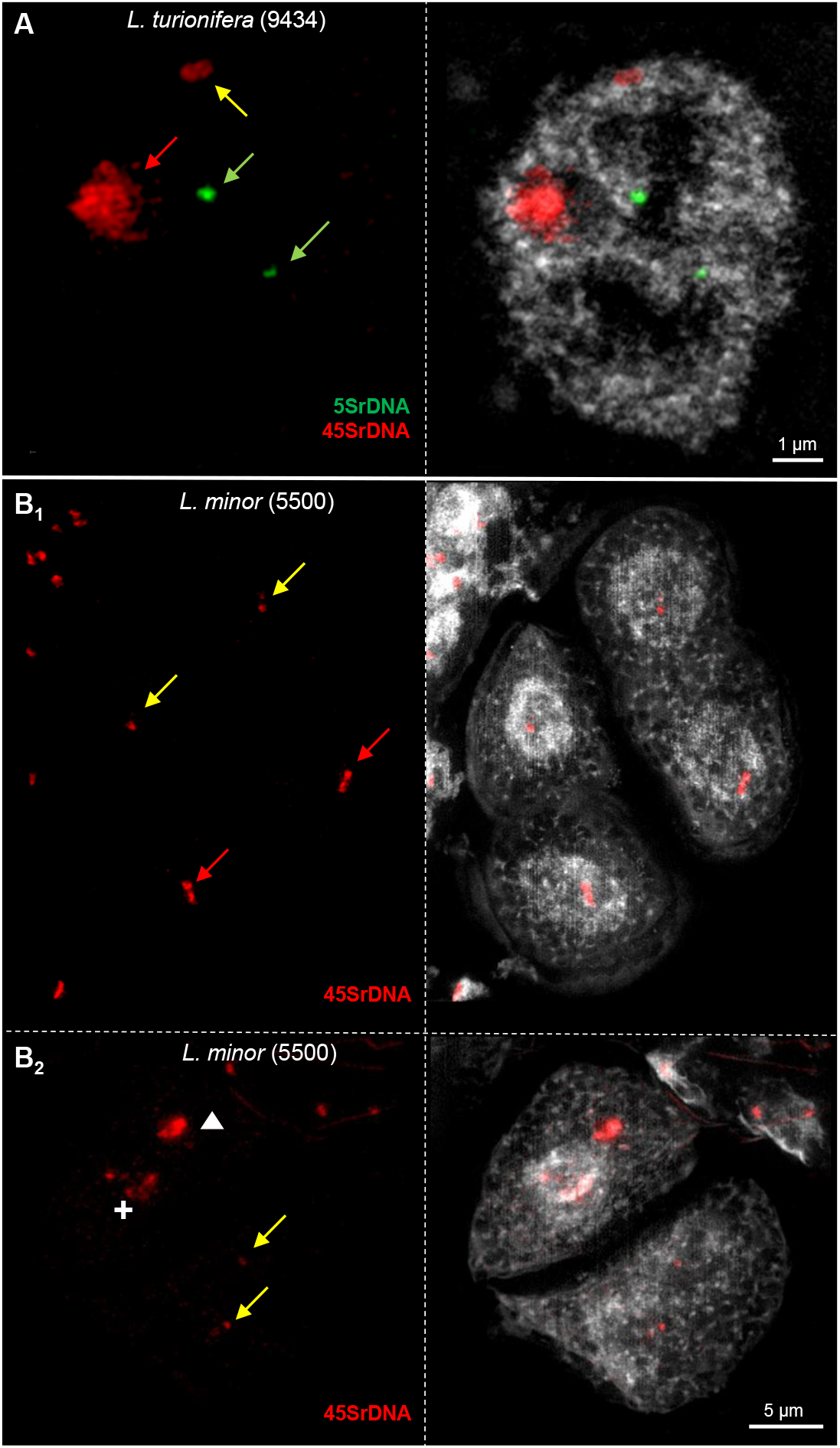
Fluorescence in situ hybridization (FISH) with rDNA detects unreduced gametes in genus *Lemna*. (A) A single slice of a 3D-SIM image stack of a somatic interphase nucleus of *L. turionifera* accession 9434 showing an extended major 45S rDNA signal attached to a nucleolus (red arrow) and a minor signal at the etch of another nucleolus (yellow arrow). Both homologous 5S rDNA signals are marked by green arrows. (B1) Minor 45S rDNA signals (yellow arrows) are visible in the two nuclei of the top cells of a tetrad. Major signals (red arrows) are in the bottom cells. (Copy number heterozygosity between homologous rDNA loci was also documented for *Spirodela polyrhiza* 5S rDNA by Stepanenko et al. (2026)). (B2) One of the major 45SrDNA signals in an unreduced dyad is dispersed within the nucleolus of the top cell (cross), the second homologous locus appears condensed outside (triangle). The bottom cell contains the two minor signals of the second homologue. The small cells adjacent to the dyad are part of the tapetum cells. Both images in (B) are maximum intensity projections, see also Supplemental movies S1 and S2.

## Discussion

Unreduced gametes, as the most important source of polyploids (followed by early somatic chromosome doubling of allodiploid zygotes), can arise through multiple cytological pathways. In plants, defective synapsis, abnormal spindle orientation, omission of meiosis I and/or II, impaired cytokinesis and restitution of nuclei were reported as causes for unreduced gametes (Bretagnolle and Thompson 1995; De Storme and Geelen 2011; Peloquin et al. 1989). Consistent with these mechanisms, our observations revealed restitution of nuclei after lacking cytokinesis during male meiosis II in *L. minor* and *L. turionifera*. These findings are the first direct evidence showing the proportions of defective products of meiosis for two *Lemna* species.

Studies in *Arabidopsis thaliana* (L.) Heynh. have identified several core meiotic regulators, such as Arabidopsis thaliana Parallel Spindle 1 (AtPS1), JASON (JAS), Omission of Second Division 1 (OSD1), Cyclin A (CYCA1;2/TAM), SUMO E3 ligase (AtMMS21) and apparently also Structural Maintenance of Chromosome 5/6 (SMC5/6), which, when disrupted, deleted or downregulated, may cause meiotic defects and result in high frequencies of unreduced gametes (d’Erfurth et al. 2008, 2010; De Storme and Geelen 2011; Liu et al. 2014; Yang et al. 2021). Based on genomic sequences, Ernst et al. (2025) proposed that *L. turionifera* and *L. minor* accessions might exhibit elevated unreduced male gamete production due to deletions in JASON homologs which are responsible for forming organelle bands and correct cytokinesis during male meiosis II (Brownfield et al. 2015). Our cytological data revealed marked inter-individual variation within this accession, with unreduced gamete frequencies ranging from 0 to 12% between individuals. Because this is substantially lower than reported for Arabidopsis mutants, a mutation of JASON with much lower penetrance could explain the observed variation in *Lemna*. OSD and TAM mutations are less likely the reason for the observed unreduced meiotic products, because fusion of dyad nuclei indicates that at least division of nuclei during meiosis II was not omitted. Mutations of SMC5/6 components could also mediate formation of unreduced gametes (Yang et al. 2021), albeit it remains unclear how such mutations disturb preferentially meiosis II, because SMC5/6 is mainly required for meiosis I.

In wild-type angiosperms lacking obvious mutations in core meiotic regulators, spontaneous unreduced gamete formation is generally rare, typically below 2% in natural populations (Kreiner et al. 2017; Ramsey 2007). The *Lemna* individuals examined here showed pronounced variation in unreduced gamete frequencies among individuals of the same clonal accessions between 0 and 12% (Table 1). While most individuals produced unreduced gametes at lower frequencies (∼1–5%), a single one of *L. minor* yielded 12% (27% of male meiotic end-stages were dyads and triads). Given the rapid clonal propagation characteristic of *Lemna*, it is possible that somatic mutations affecting meiotic stability were acquired independently among and within different clonal accessions, contributing to this heterogeneity. As an alternative to mutation of meiotic genes, environmental stress causing a switch to sexual propagation including emergence of a few unreduced gametes could be responsible for natural auto- and allopolyploids. In vitro flowering requires hormonal treatments that mimic natural stress responses, and such stress has been linked to increased frequencies of unreduced gametes in plants (Ramsey and Schemske 1998). Under unfavorable conditions that constrain asexual propagation, sexual reproduction could therefore contribute to adaptive genetic diversity, potentially through polyploidization and hybrid formation (Mason and Pires 2015).

Although not easily accessible, unreduced *L. minor* female gametes should also occur, since triploid *L*.*× japonica* with two genomes of *L. minor* were found, albeit crossing between of *L. minor × L. turionifera* was exclusively unidirectional as was shown by plastid markers (Braglia et al. 2021; Ernst et al. 2025).

Despite their predominantly clonal reproduction, *L. minor* and *L. turionifera* conform to general patterns reported across angiosperms (Bretagnolle 2001; Ramsey 2007), in which a higher frequency of unreduced gametes is restricted to relatively few individuals. Notably, tetraploid *L*. × *japonica* has not been reported so far. Low flowering rates of the parental species in natural populations (1.5–5%; Landolt 1986, pp. 167□169), together with low frequencies of unreduced gametes, substantially reduce the probability of fusion of unreduced gametes from both parents. This explanation may also account for the absence of autotetraploid *L. minor* and *L. turionifera*.

Our study provides a fundament for comparative cytological analyses of meiotic stages in duckweeds. Diverse natural allopolyploids are even more common in sect. *Alatae* (Stepanenko et al. 2025) than in sect. *Lemna*, exemplified by the fertile tetraploid taxon *L*. × *aoukikusa* Beppu et Murata and its triploid backcrosses with *L. aequinoctialis*. Most likely, interspecific hybrids occur also in the genus *Wolffia* Horkel ex Schleid. Thus, knowledge on how to address meiotic processes (this paper) and on species fertility and crossability (Lee et al. 2025) will uncover lineage-specific routes underlying polyploidization and diversification in duckweeds and provide prerequisites to harness crossing and thus domestication of economically interesting duckweeds.

## Materials and methods

### Plant materials and culture conditions

All tested duckweed accessions were originally from the Landolt collection and are now available at the duckweed collection of the Institute of Agricultural Biology and Biotechnology (CNR-IBBA) in Milan (Italy; Morello et al. 2025). The diploid accessions *L. minor* 5500 and *L. turionifera* 9434 and 7432 were selected based on prior investigations of their floral development as well as their genetic and cytogenetic features (Braglia et al. 2021; Ernst et al. 2025; Lee et al. 2025). Plants were maintained under axenic conditions, see Braglia et al. (2021).

### Preparation of meiotic cells

To prepare meiotic stages, young developing anthers were collected from artificially induced flowers (Lee et al. 2025). In brief, few fronds were precultured for seven days in 50 ml plastic beakers with 20 ml of SH medium (Schenk and Hildebrandt 1969) including 1% sucrose. For flower induction (FI), some asexually propagating, healthy fronds were then put in 100 ml plastic beakers with 100 ml of modified NH_4_^+^ free 5% Hutner liquid medium (Lee et al. 2025) and 30 µM salicylic acid (SA). The plants were kept under a 16 h light/8 h dark photoperiod (white light, 150 μmol mD^□2^ sD^□1^) at 25□. Considering that *Lemna* plants usually start flowering from day 12–14 of FI (Lee et al. 2025), at day 9, fronds were transferred into 90 × 20 mm petri dishes containing the same FI medium made semisolid by 0.3% agar, to monitor for expected immature flowers (possibly undergoing meiosis).

Fronds bearing young flowers were confirmed by checking the underside of the petri dish under a stereo microscope (SMA18, Nikon, Tokyo, Japan). Immature flowers of different size were independently dissected and immediately fixed in a freshly prepared 3:1 (v:v) ethanol:acetic acid solution for 3–6 days at 4°C, then transferred to 70% ethanol and stored at –20□°C until use.

Prior to slide preparation, fixed flowers were treated with 45% acetic acid for up to 30 min. Single anthers were transferred onto glass slides with a drop of 1 µl 45% acetic acid and further dissected by hypodermic needles to release the meiotic cells. After adding 7 µl of 45% acetic acid, the cell suspension was covered with a coverslip. Slides were briefly flamed over burning ethanol (∼1 sec), then kept in liquid nitrogen for 10 sec. The coverslip was subsequently removed using a razor blade, and the slides were immediately placed in pre-chilled 99.8% ethanol at –20°C.

### Probe generation and fluorescence in situ hybridization (FISH)

*Lemna-*specific 5S and 45S rDNA probes were generated following the procedure described by Hoang and Schubert (2017) and Stepanenko et al. (2026) with small modifications. First, genomic DNA isolated from *L. minor* (clone 8703) was amplified using duckweed rDNA-specific primers (Hoang and Schubert (2017); Stepanenko et al. (2026)). After purification, the obtained PCR products were used as templates for labeling with specific fluorescent dyes. The rDNA fragments were subjected to nick-translation using the AZDye594 NT Labeling Kit for 45S rDNA (red) and the AZDye488 NT Labeling Kit for 5S rDNA (green) (Jena Bioscience, Jena, Germany). After precipitation with 96% ethanol, probe pellets were dissolved in 100 μl hybridization buffer (50% (v/v) formamide, 20% (w/v) dextran sulfate in 2 × SSC, pH 7).

The meiotic products prepared on slides were denatured in 2N NaOH for 5 min at room temperature (RT), rinsed, dehydrated in 75% (twice) and 99.8% ethanol for 5 min each, and air-dried. Of each FISH probe, 1 µl (50 ng/μl) was added to 10 μl hybridization mixture [dextran sulphate (1 g) dissolved in 1 ml ddH□O, followed by addition of 5 ml deionized formamide, 1 ml 20 × SSC, and 1 ml sonicated fish sperm DNA], denatured at 99°C for 15 min, and stored at −20□°C until use. The mixture was applied to air-dried slides, sealed with coverslips, and incubated in a moist chamber at 37□°C overnight (Aliyeva-Schnorr et al. 2015). After hybridization, the slides were rinsed twice in 2 × SSC for 5□min, followed by distilled water at RT for 2□min, and then air-dried. Finally, 10□μl of DAPI antifade solution (1□μg/ml) was added to each slide and sealed with a coverslip.

### Microscopy

To detect the ultrastructural chromatin organization of cells at a resolution of ∼120 nm (super-resolution achieved with a 488 nm laser excitation), spatial structured illumination microscopy (3D-SIM) was performed with a 63×/1.4 Oil Plan-Apochromat objective of an Elyra 7 microscope system and the software ZENBlack (Carl Zeiss GmbH). Images were captured separately for each fluorochrome using the 405, 488, and 561 nm laser lines for excitation and appropriate emission filters (Weisshart et al. 2016). The spatial animation (Movie S2) was done based on the SIM image stack (Movie S2) using the Imaris 9.7 (Bitplane) software. The tool ‘Surface’ was applied to indicate the nuclei volumes. Microspores stained with DAPI were counted using the fluorescence microscope AXIO Lab.A1 (Carl Zeiss GmbH).

## Acknowledgements

We appreciate our colleagues in both IBBA and IPK for their kind support, in particular Dr. Stefan Heckmann and Dr. Ewa Piskorz (both from IPK) for critical reading of the manuscript. Anton Stepanenko was supported by a Ukraine Distinguished Fellowship of the German National Academy of Sciences Leopoldina. This study was carried out within project AGRITECH National Research Center funded by EU Next-Generation EU PNRR Missione 4 Componente 2, Investimento 1.4 – (D.D. 1032 17/06/2022, CN00000022) to IBBA-CNR, and Project IR0000032 – ITINERIS - Italian Integrated Environmental Research Infrastructures System. We acknowledge IBISBA-IT CNR-IBBA node for the access provided to the infrastructure (microscope). This manuscript reflects only the authors’ views and opinions, neither the European Union nor the European Commission can be considered responsible for them.

## Author contribution

YL, IS and LM designed the research. YL, VS, AS, GK and LB performed research. IS and LM supervised. YL wrote the original manuscript. IS and LM edited the manuscript. LM and LB administrated project & funding.

## Data availability statement

All data are reported in the manuscript.

## Material distribution footnote

The author responsible for distribution of materials integral to the findings presented in this article in accordance with the policy described in the Instructions for Authors (https://academic.oup.com/plcell/pages/General-Instructions) is: Laura Morello

## Publication ethics

The authors declare no competing interests.

---

**Suppl. figure 1.**
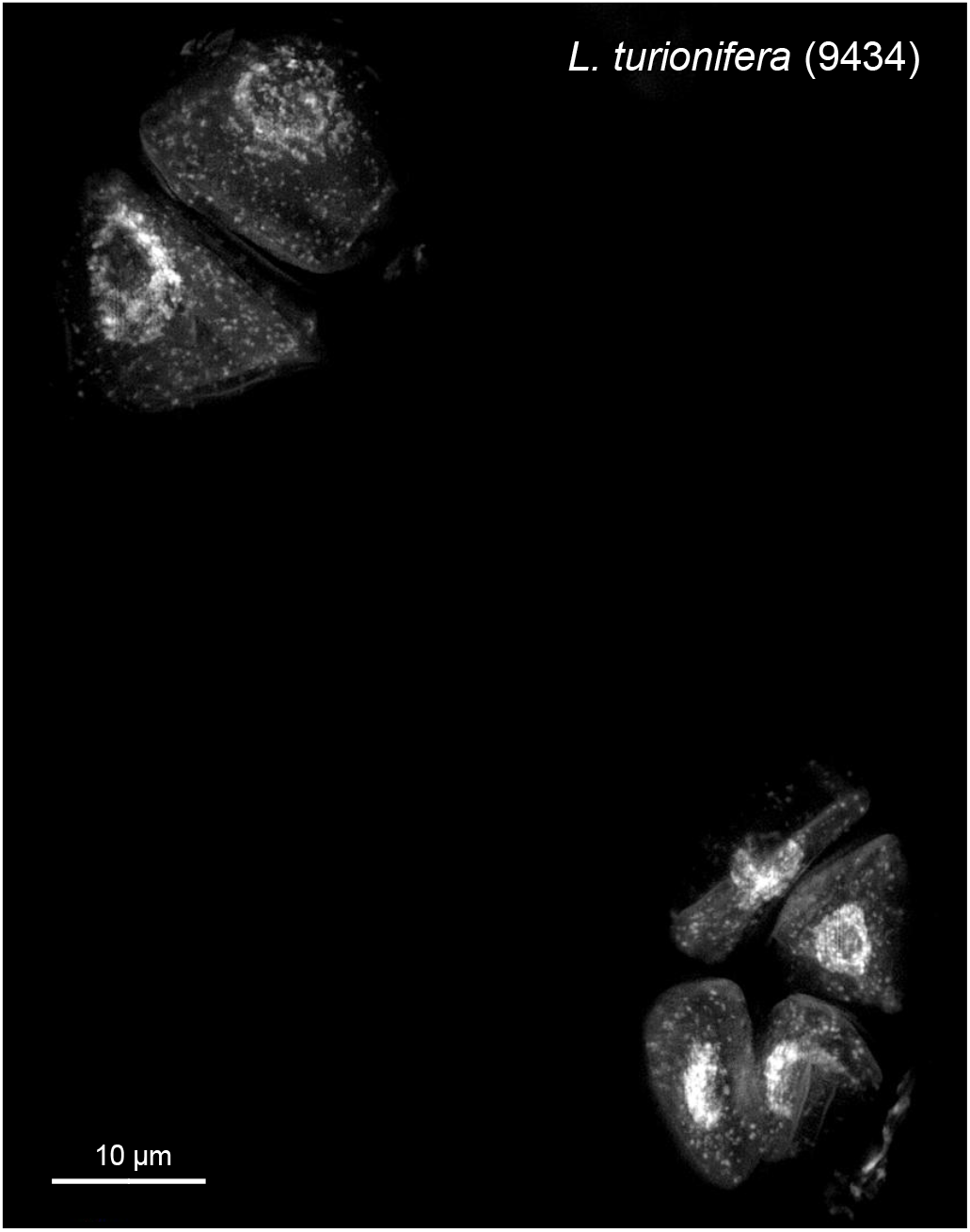
Dyad (top) and tetrad of *Lemna turionifera* (9434).

**Suppl. Movie 1.**
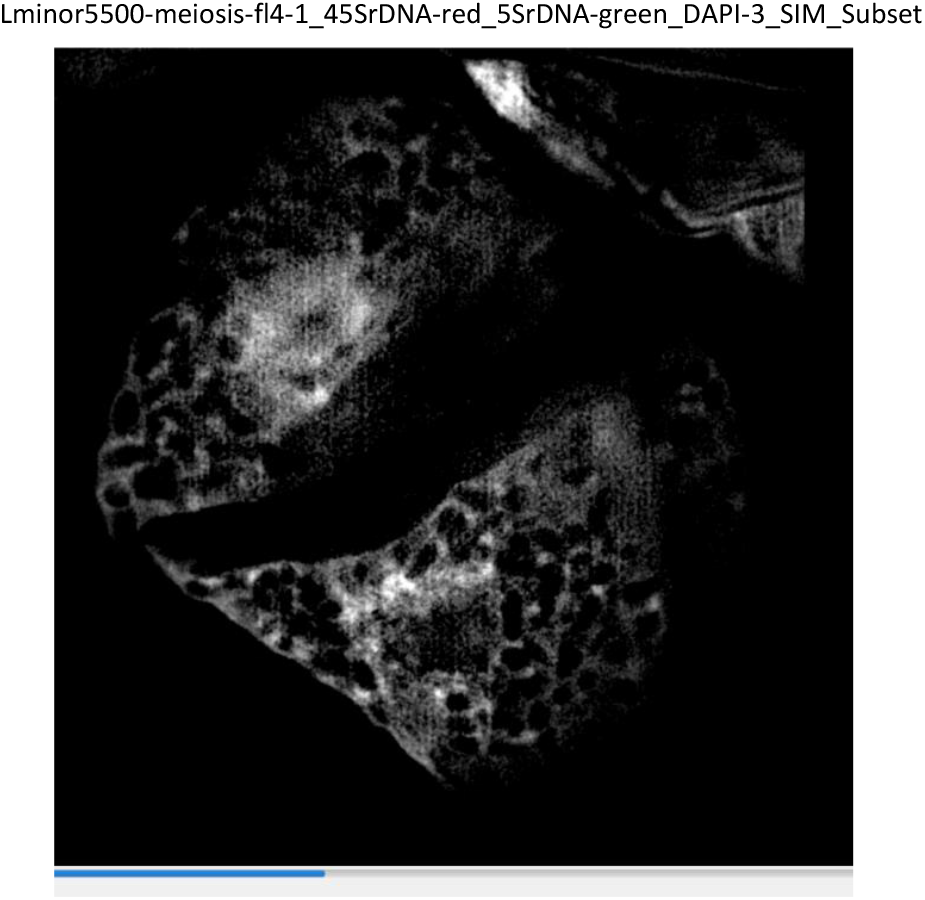
Image stack of the *Lemna minor* (5500) dyad shown in fig.3B_2_ and Suppl. Movie 2.

**Suppl. Movie 2.**
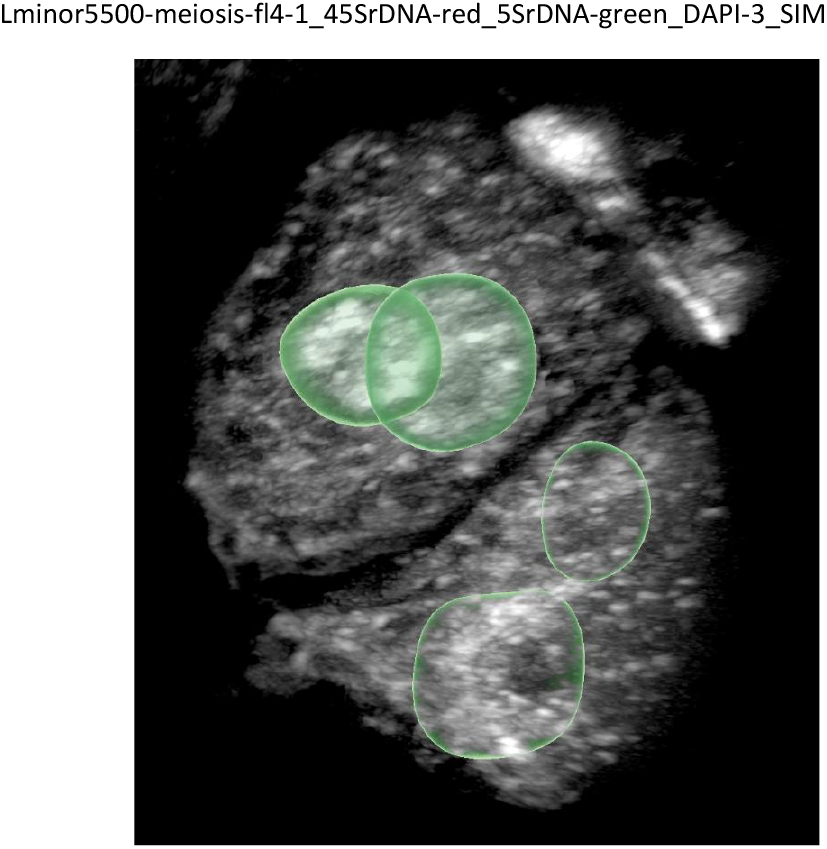
3D animation of the *Lemna minor* (5500) dyad shown in fig.3B_2_ and Suppl. Movie 1. The green spheres indicate the nuclei in both cells.

**Optional suppl. fig.**
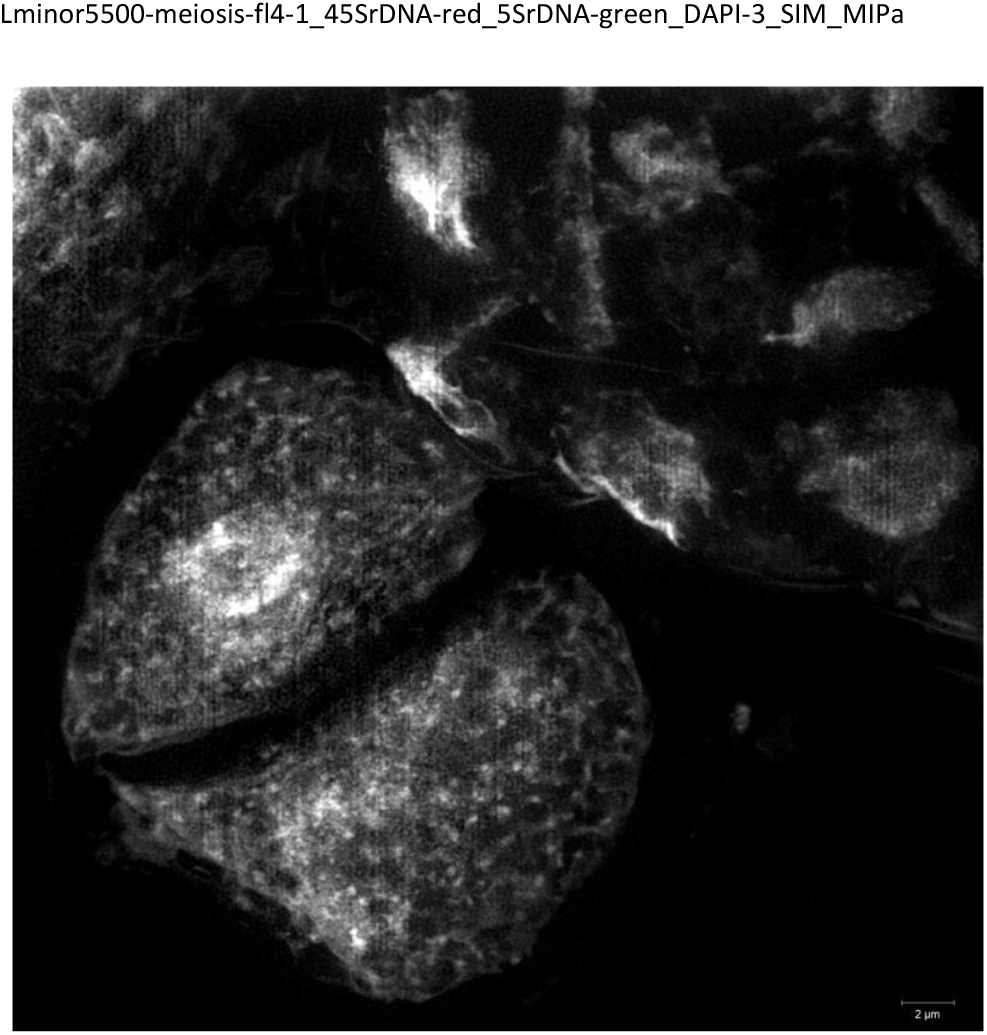
fig. Extended fig.3B_2_ and of Suppl. Movies 1 and 2.

## Notes

### Competing Interest Statement

The authors have declared no competing interest.

